## Supplementary text for "Leveraging functional annotation to identify genes associated with complex diseases"

Supplementary methods

**Variational Bayes model for variable selection in imputing gene expression**

Prior assumptions on hyper parameters

In practice, we do not have any prior information to select the values of $a,b,c,d,a_{0}$ and $b_{0}$, especially for
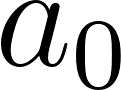
 and
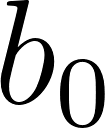
. Therefore, twenty grid points of $\pi$ are initially sampled from the following distribution, as what have been proposed in the literature [1, 2]:

$$\log\left( \frac{\pi_{k}}{1-\pi_{k}} \right)\sim unif(-\log_{10} \left( p \right), -1)$$

*Updating beta coefficients*:

For updating the SNP coefficient $\beta_{k}$ of the SNP $k$ in the model, we would get:

$$\ln Q^{*}\left( \beta_{k} \right)=E_{rest}\left[ \ln P\left( Y | \beta,\epsilon,X \right)+\ln P\left( \beta| \sigma_{\beta}^{2},\sigma^{2},\gamma\right) \right]+constant$$

$$=E_{rest}\left[ \sum_{i=1}^{n} -\frac{\left( y_{i}-x_{i}\beta\right)^{2}}{2\sigma^{2}}+\sum_{j=1}^{p} -\frac{\gamma_{j}\beta_{j}^{2}}{2\sigma_{\beta,j}^{2}\sigma^{2}} \right]+constant$$

$$=E_{rest}\left[ \sum_{i=1}^{n} -\frac{\left( y_{i}-x_{i}\beta\right)^{2}}{2\sigma^{2}}-\frac{\gamma_{k}\beta_{k}^{2}}{2\sigma_{\beta,k}^{2}\sigma^{2}} \right]+constant$$

Therefore, by taking the expectation of the other parameters, $\beta_{k}$ approximately follows a normal distribution where:

$$s_{k}^{2}=Var\left( \beta_{k} | \gamma_{k}=1 \right)= \frac{\sigma^{2}}{\left( X^{T}X \right)_{kk}+1/\sigma_{\beta}^{2}}$$

$$\mu_{k}=E\left( \beta_{k} | \gamma_{k}=1 \right)=\frac{s_{k}^{2}}{\sigma_{k}^{2}}[\left( X^{T}X \right)_{k}-\sum_{j\neq k} \left( X^{T}X \right)_{jk}\alpha_{j}\mu_{j}]$$

$$\alpha_{k}=P\left( \gamma_{k} =1 \right|rest)$$

*Updating the probability of being selected*

The probability of the kth SNP getting selected is dependent not only on the relationship between the SNP genotype and phenotype (gene expression level here), but also on the epigenetic signals of the SNP, which can be described through the equation below:

$$\ln Q^{*}\left( \gamma_{k} \right)=E_{rest}\left[ \ln P\left( Y | X,\beta,\sigma^{2} \right)+\ln P\left( \beta| \gamma,\sigma_{\beta}^{2},\sigma^{2} \right)+\ln P\left( \gamma| A,\omega\right) \right]+constant$$

After solving the equation above term by term, we have:

$$\ln Q^{*}\left( \gamma_{k}=1 \right)=\frac{\mu_{k}^{2}}{2s_{k}^{2}}+\frac{1}{2}\ln\sigma^{2}\sigma_{\beta}^{2}+\ln\frac{1}{1+\exp\left( -A_{k}\omega\right)}+constant$$

$$\ln Q^{*}\left( \gamma_{k}=0 \right)=\ln\frac{1}{1+\exp\left( A_{k}\omega\right)}+constant$$

$$\frac{\alpha_{k}}{1-\alpha_{k}}=\exp\left( \frac{\mu_{k}^{2}}{2s_{k}^{2}} \right)\times\frac{1}{\sigma_{\beta}^{2}\sigma^{2}}\times\frac{1+\exp(A_{k}\omega)}{1+\exp({-A}_{k}\omega)}$$

*Updating coefficients in the logit link of epigenetics signals*

Parameters in the logit link are related to both the SNP epigenetic signals and the status of the SNP being selected or not in last updating step which shown in the following:

$$\ln Q^{*}\left( \omega\right)=E_{rest}\left[ \ln P\left( \gamma| A,\omega\right)+\ln P\left( \omega| \eta\right) \right]+constant$$

Similar to the idea of using variational Bayes method in a logistic regression introduced in the literature [3], $\omega$ also has a normal distribution with parameters:

$$\omega_{N}=E\left( \omega| rest \right)=V_{N}\sum_{j=1}^{p} \frac{\gamma_{j}}{2}A_{j}^{T}$$

$$V_{N}^{-1}=\frac{1}{Var(\omega|rest)}=E_{rest}\left( \eta\right)I+2\sum_{j=1}^{p} \lambda\left( \xi_{j} \right)A_{j}^{T}A_{j}$$

To calculate the overall PPS, we need to get the likelihood of the full model under each prior parameter setting. Since we actually approximate the largest value of the lower bound in the full model likelihood, the estimated optimal lower bound is used to substitute for the exact likelihood.

$$\ln P\left( Y | X,Q,A \right)\geq F\left( Q,\theta\right)$$

$$=\int\int\int q(\beta,\gamma,\omega;\theta)log \frac{P(Y,\beta,\gamma,\omega|X,A;\theta)}{q(\beta,\gamma, \omega;\theta)}d\beta d\gamma d\omega$$

$=E_{\beta,\gamma, \omega}\left[ \log P\left( Y,\beta,\gamma,\omega| X,A;\theta\right) \right]-E_{\beta,\gamma, \omega}[\log q(\beta,\gamma,\omega|\theta)]$

where

$$\log P\left( Y,\beta, \gamma,\omega| X,A,\theta\right)=\log P\left( Y | \beta,\sigma^{2} \right)+\log P(\beta|\gamma,\sigma^{2},\sigma_{\beta}^{2})+\log P\left( \gamma| \omega,A \right)+\log P(\omega|\eta)$$

$$\log q\left( \beta,\gamma,\omega;\theta\right)=\log q\left( \beta| \gamma;\theta\right)+\log q\left( \gamma| \omega;\theta\right)+\log q(\omega;\theta)$$

Then, the lower bound becomes:

$$F\left( Q,\theta\right)=-\frac{n}{2}\log2\pi\sigma^{2}-\frac{\left| \left| Y-X\alpha\mu\right| \right|^{2}}{2\sigma^{2}}-\frac{1}{2\sigma^{2}}\sum_{j=1}^{P} [\left( X^{T}X \right)_{jj}Var(\beta_{j})]$$

$$-\sum_{j=1}^{P} \alpha_{j}\log\frac{\alpha_{j}}{\pi}-\sum_{j=1}^{P} \left( 1-\alpha_{j} \right)\log\frac{1-\alpha_{j}}{1-\pi}$$

$$+\sum_{j=1}^{p} \frac{\alpha_{j}}{2}(1+\log\frac{s_{k}^{2}}{\sigma^{2}\sigma_{\beta}^{2}}-\frac{s_{k}^{2}+\mu_{k}^{2}}{\sigma^{2}\sigma_{\beta}^{2}})$$

$$+\frac{1}{2}w_{N}^{T}v_{N}^{-1}w_{N}+\frac{1}{2}\ln\left| V_{N} \right|+\sum_{j=1}^{P} [-\ln\sigma\left( \xi_{j} \right)-\frac{\xi_{j}}{2}+\lambda\left( \xi_{j} \right)\xi_{j}^{2}]$$

$$-\ln\Gamma\left( a_{0} \right)+a_{0}\ln b_{0}-b_{0}\frac{a_{N}}{b_{N}}+\ln\Gamma\left( a_{N} \right)+a_{N}$$

The implementation of the method utilized part of the script used in the varbvs R package [1]. The running time of our method is longer compared to that of elastic net (glmnet R package) [4] when fitting models on a data set with 100 individuals, 1,000 SNPs and 5 kinds of annotation array. Our method takes 15.15 minutes while fitting models repeated for 100 times while elastic net takes 13.90 seconds on the same CPU (Intel(R) Xeon(R) CPU E5-2620_v3 2.40GHz).

Supplementary Discussion

**Gene expression imputation accuracy and gene-trait association test**

Gene expression imputation accuracy was compared across all five methods through both five-fold cross validation and prediction in an external dataset from the CommonMind Consortium ([www.synapse.org/CMC](http://www.synapse.org/CMC)). We compared the $R^{2}$ (squared correlation) between the observed and imputed expression levels of our method with that from the other methods. In five-fold cross-validation analysis using the GTEx data (**S5 Fig**), our method showed $R^{2}$ decrease from 6.3% (-2.4e-3,compared with vb.annot) to 22.06% (7.2e-3,compared with elastic net), compared to the other four methods (Fig 5, supplementary). For prediction analysis in the CommonMind dataset, expression imputation models trained on the GTEx Brain Cortex BA9 tissue were used to predict gene expression levels based on individual genotype data of CommonMind. All five methods had similar $R^{2}$ (~0.007) between predicted gene expression and whole expression in the Brain Cortex BA9 tissue of CommonMind (**S6 Fig**) but we note that our method included more genes whose expression levels can be predicted. For all five methods, in-sample cross-validation performance was much better than that in out-sample prediction as expected.

We further investigate the potential influence of imputation accuracy on final gene-trait association analysis. No consistent nor significant associations between imputation accuracy and the percentage of associated genes having pLI>0.99 were identified in the five methods (**S7 Fig**). Similarly, the percentage of genes identified as trait-associated across 208 traits has no consistent association between imputation accuracy across five methods (**S8 Fig**).
